# Benchmarking Uncertainty and Improving Risk Ranking in Multi-task Bioactivity Prediction

**DOI:** 10.64898/2026.09.15.751772

**Authors:** Zijian Wang, Li Tian, Chi-Ming Che

## Abstract

Multi-task bioactivity prediction transfers information across assays, but experimental prioritization also requires uncertainty estimates that identify unreliable predictions. We benchmarked prediction accuracy and uncertainty calibration for five multi-task predictors on 100 ChEMBL27 test assays and 100 CHEMBL37 assays under the random split and the clustering split. Calibration used native, Deep Ensemble and MC Dropout uncertainty; risk ranking was compared for two Gaussian process (GP) backbones. Adaptive deep kernel fitting (ADKF) achieved the best mean calibration results, while deep kernel transfer (DKT) ranked errors more effectively than ADKF with native uncertainty. We also introduce Influence Calibrated Support Reconstruction (ICSR), a risk score for kernel-based predictors. ICSR combines measured support reconstruction errors with query-specific influence and stabilizes their weighted average toward the full-support mean to adjust Gaussian process uncertainty while preserving predicted activities. With DKT and ADKF, ICSR improved all three mean risk-ranking metrics over native uncertainty and an adapted neighborhood comparator in every panel and split. ICSR achieved a mean half-query MAE reduction (*R*_50_) of 14.2–19.6%, where *R*_50_ measures the percentage decrease in mean absolute error (MAE) after retaining the lowest-risk half of the queries. A retrospectively selected SARS-CoV-2 main protease case further illustrated its use for selecting more reliable predictions. The benchmark supports joint assessment of calibration and error ranking, while ICSR improves selective use of kernel-based bioactivity predictions.

## 1. Introduction

Multi-task bioactivity prediction transfers structure–activity information across assays to support prediction from limited measurements.^1,2^ Each assay defines a task. At evaluation, measured compounds form the support set used to condition or adapt the predictor, while query activities are withheld for assessment. One-shot molecular learning demonstrated the value of across-assay transfer,^3^ whose benefit depends on the chemistry and targets represented in the training assays.^4^ Experimental prioritization also requires identifying which predictions are reliable within the assay of interest.

A benchmark of uncertainty quantification (UQ) in this setting must distinguish prediction accuracy, calibration and risk ranking: small average errors, uncertainty scales that match observed errors, and effective ordering of queries by error risk are different goals. Molecular UQ benchmarks have shown that performance depends on the predictor, data and evaluation objective.^5–8^ Recent studies have compared uncertainty estimators in proteochemometric bioactivity models^9^ and multi-task molecular property models under data shifts.^10^ These studies motivate a focused question for assay transfer: how do the uncertainty estimators associated with different predictors compare on the same assays and support–query splits?

A second question is whether measured supports can improve query risk assessment beyond the uncertainty returned by a predictor. Deep kernel learning couples learned representations to Gaussian process (GP) covariance functions.^11^ Deep kernel transfer (DKT) and adaptive deep kernel fitting (ADKF) use this framework to combine representations learned across tasks with GP prediction within each assay.^12,13^ Their native predictive standard deviations (SDs) are conditional on the fitted representation and covariance model^14^ and may miss local discrepancies between predictions and measured activities. Leave-one-out (LOO) reconstruction examines these discrepancies by predicting each measured support from the others, with analytic GP identities available when the fitted model is held fixed.^14,15^

LOO errors have been used for GP hyperparameter selection^16^ and model diagnostics,^17^ while standardized squared LOO errors can estimate an overall variance scale under covariance misspecification.^18^ Local precedents include DPRESS, which redistributes squared cross-validation errors through descriptor-space distances,^19^ and the different-neighbor-ratio GP (GP-DNR), which supplements GP uncertainty with predicted activity variation among structurally similar compounds.^20^ Expected squared LOO errors have also guided adaptive sampling with GP emulators.^21^ These approaches motivate using measured reconstruction errors locally; the methodological question here is how to assign those errors to queries according to the fitted predictor’s dependence on each support.

We address these questions through two contributions. First, we benchmark prediction accuracy and uncertainty calibration for DKT, ADKF, conditional neural processes (CNP), ActFound and MetaNN on 100 ChEMBL27 test assays and 100 CHEMBL37 assays under the random split and the clustering split. The comparison covers native model-output uncertainty, Deep Ensembles and MC Dropout as applicable to these predictors. Shared support–query splits allow prediction accuracy and calibration to be assessed under the same assay conditions.

Second, we introduce Influence Calibrated Support Reconstruction (ICSR), a risk score for kernel-based multi-task bioactivity prediction (Figure 1). ICSR assigns standardized squared support reconstruction errors to each query through influence weights defined by the fitted prediction’s sensitivity to measured support activities. Stabilizing the weighted errors toward their full-support mean gives a query-specific adjustment to native GP uncertainty while preserving predicted activities. Matched comparisons with native SD and an adapted GP-DNR on DKT and ADKF test the contribution of this score to risk ranking and selective prediction. Reconstruction controls examine the information supplied by correctly located support errors, and a retrospective SARS-CoV-2 main protease (Mpro) case illustrates selection within a chemical series containing close query–support structural analogues.

**Figure 1.**
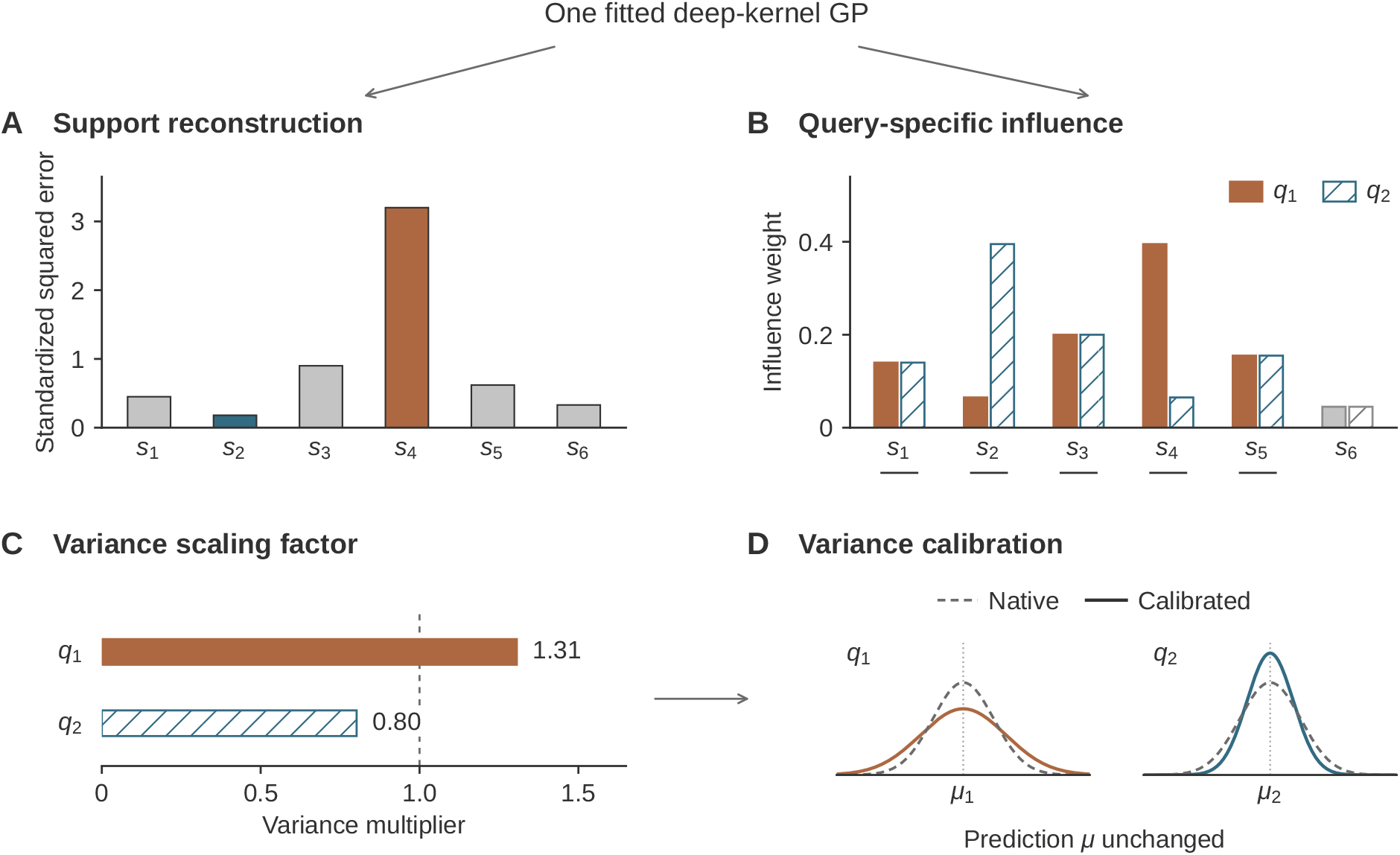
ICSR calibrates predictive variance while preserving predicted activity *μ*. Supports (*s*) are measured compounds; queries (*q*) are compounds to be predicted. (A) Each support is reconstructed from the others; its squared error is divided by its conditional variance. Both queries share these standardized errors. (B) *q*_1_ emphasizes poorly reconstructed *s*_4_; *q*_2_ emphasizes well-reconstructed *s*_2_. Swapping the weights of *s*_2_ and *s*_4_ preserves weight concentration. Filled and hatched bars identify *q*_1_ and *q*_2_, respectively. Underlined supports enter local weighting; gray *s*_6_ bars are excluded. All six supports remain in GP conditioning and reconstruction. (C) Influence-weighted errors, stabilized toward the full-support mean, give variance multipliers of 1.31 and 0.80. The dashed line at one denotes unchanged variance. (D) Schematic Gaussian profiles widen or narrow around unchanged *μ*_*q*_ values (dashed, native; solid, calibrated); width changes are exaggerated for visual clarity. Values in A–C are illustrative, using *η* = 0.75 and DKT stabilization *κ* = 4. ICSR is the square root of the scaled variance.

## 2. Results

All predictors evaluated in this section were trained on the same 4,100 assays from the ChEMBL27 collection, which contains 4,276 assays prepared through the profile QSAR (pQSAR) workflow.^1^ Of the remaining assays, 76 were used for validation and hyperparameter tuning and 100 were held out for internal testing. To assess generalization beyond this collection, we applied the pQSAR authors’ public extraction and filtering pipeline to a later ChEMBL release, CHEMBL37. After excluding all 4,276 ChEMBL27 assay identities, we uniformly sampled 100 assays without replacement from 4,261 eligible assays for external testing.

Across these two 100-assay test sets, we first benchmarked prediction accuracy and uncertainty calibration for five multi-task predictors and their associated uncertainty estimators, using shared support–query splits under the random split and the clustering split. We then compared native SD, GP-DNR and ICSR for risk ranking and selective prediction with DKT and ADKF, keeping predictions identical across scores within each fitted model. Reconstruction controls and a retrospective SARS-CoV-2 Mpro application case examined the source and practical use of these gains.

### 2.1 Multi-task prediction accuracy depends on the support–query split

Mean absolute error (MAE) is the mean absolute difference between predicted and measured query activities, on the recorded assay scale. Squared Pearson correlation (*r*^2^) is the square of their Pearson correlation and measures the strength of unsigned linear association. Lower MAE and higher *r*^2^ are favorable. We used these two metrics to assess DKT, ADKF, CNP, ActFound and MetaNN (Figure 2). The ChEMBL27 and CHEMBL37 panels each contain 100 assays evaluated with the clustering split and count-matched repeats of the random split.

**Figure 2.**
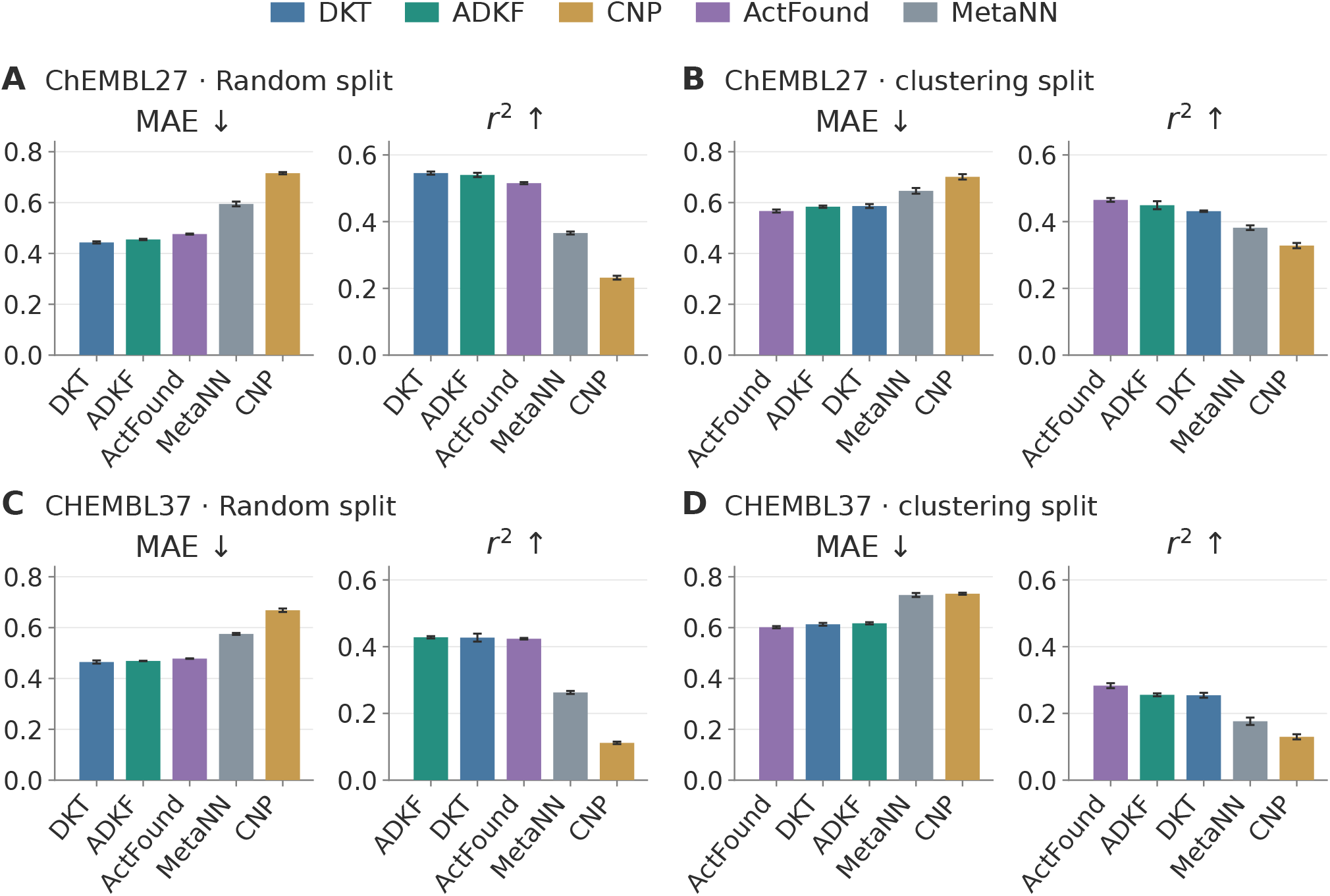
Prediction accuracy across five multi-task predictors. Mean absolute error (MAE) is the mean absolute query prediction error on the recorded assay scale; squared Pearson correlation (*r*^2^) measures unsigned linear association between predicted and measured activities. Lower MAE and higher *r*^2^ are favorable. Panels show ChEMBL27 under the random split (A) and clustering split (B), and CHEMBL37 under the random split (C) and clustering split (D), with 100 assays per dataset. Within each panel and metric, algorithms are ordered from best to worst using unrounded means: ascending MAE and descending *r*^2^. Algorithm colors are fixed across panels. Bars show means, and error bars show between-run sample SDs across four independent training runs. Metrics are calculated within each support– query split; five random repeats are averaged within assay before equal assay weighting, while the clustering split is evaluated once per assay. Each metric has its own scale, fixed across panels; all bar axes start at zero.

MAE quantifies average absolute prediction error, while *r*^2^ quantifies squared linear association; lower MAE and higher *r*^2^ are favorable. Across both split regimes, ActFound and the two kernel-based models had lower mean MAE and higher mean *r*^2^ than MetaNN and CNP (Figure 2). Their relative performance depended on the split: the random split favored DKT and ADKF, whereas the clustering split favored Act-Found in both panels. The differences among these three models were smaller than their gaps to MetaNN and CNP.

MAE measures average absolute prediction error, and *r*^2^ measures squared prediction–activity correlation; lower MAE and higher *r*^2^ are favorable. Under the random split, DKT and ADKF had lower mean MAE and slightly higher mean *r*^2^ than ActFound, but these advantages were modest. The differences were particularly small on CHEMBL37. DKT was slightly better than ADKF on ChEMBL27, while their CHEMBL37 results were nearly identical; despite its adaptive GP head, ADKF thus showed little gain over DKT in this setting. MetaNN and CNP had higher MAE and lower *r*^2^ than these three models on both panels, with MetaNN consistently outperforming CNP.

MAE expresses average absolute prediction error on the recorded assay scale; *r*^2^ expresses squared linear association. Lower MAE and higher *r*^2^ are favorable. Under the clustering split, ActFound had lower mean MAE and higher mean *r*^2^ than both kernel-based models in both panels, although the differences were modest. ADKF, which adapts its GP head to the support set, improved on DKT’s globally fitted head on both metrics on ChEMBL27, but did not close the gap to ActFound. On CHEMBL37, ADKF had slightly higher MAE and slightly higher *r*^2^ than DKT, leaving neither GP model better on both metrics. MetaNN and CNP remained less accurate than all three models; their CHEMBL37 MAE values were close, although MetaNN retained higher *r*^2^. The clustering split generally increased MAE, with CNP on ChEMBL27 providing an exception. Changes in *r*^2^ also differed across models: it increased for CNP on both panels and for MetaNN on ChEMBL27, whereas it decreased for ActFound and both GP models.

MAE is average absolute prediction error; paired MAE comparisons assess its difference between predictors evaluated on matched assay splits, with lower MAE favorable. ActFound combined the lowest mean MAE under the clustering split with competitive MAE under the random split. Targeted paired MAE comparisons supported its advantage under the clustering split over DKT only on ChEMBL27, with an interval narrowly excluding zero, and over ADKF only on CHEMBL37 (Table 2C). Under the random split, both GP models’ MAE advantages over ActFound reached significance on ChEMBL27, but neither CHEMBL37 comparison did. Only part of the mean MAE ordering was therefore supported statistically.

### 2.2 Native uncertainty estimates are better calibrated than ensemble and dropout estimates

We used MCA, CE90, NLL and ENCE to evaluate uncertainty calibration for DKT, ADKF, CNP, ActFound and MetaNN (Table 1). Nominal interval coverage is a prediction interval’s target probability; observed interval coverage is the fraction of all query activities inside their corresponding intervals in one support– query split. Miscalibration area (MCA) integrates the absolute difference between observed and nominal interval coverage across nominal levels. Absolute interval coverage error at 90% (CE90) is the absolute difference at the nominal 90% level, computed per split before averaging and expressed in percentage points. Gaussian negative log likelihood (NLL) is the mean negative natural logarithm of the Gaussian density assigned to measured query activities; it assesses both prediction error and uncertainty width. Expected normalized calibration error (ENCE) averages |RMSE−RMV|/RMV across ten bins ordered by predictive uncertainty, where root mean squared error (RMSE) is the square root of mean squared prediction error and root mean predicted variance (RMV) is the square root of mean predicted variance within the same bin. Lower values are favorable for all four metrics.

**Table 1.** Calibration of native, Deep Ensemble and MC Dropout uncertainty.

| Score | Model | ChEMBL27 (100 assays) |  |  |  | ChEMBL37 (100 assays) |  |  |  |
| --- | --- | --- | --- | --- | --- | --- | --- | --- | --- |
|  |  | MCA ↓ | CE90 ↓ | NLL ↓ | ENCE ↓ | MCA ↓ | CE90 ↓ | NLL ↓ | ENCE ↓ |
| *Random split* |  |  |  |  |  |  |  |  |  |
| Native SD | DKT | **0.139 ± 0.012** | **15.9 ± 2.6** | **1.463 ± 0.153** | **0.733 ± 0.076** | **0.166 ± 0.022** | **24.1 ± 4.5** | **1.832 ± 0.333** | **0.908 ± 0.139** |
| Native SD | ADKF | **0.101 ± 0.001** | **6.2 ± 0.1** | **1.006 ± 0.004** | **0.471 ± 0.001** | **0.077 ± 0.002** | **5.8 ± 0.1** | **0.891 ± 0.005** | **0.422 ± 0.003** |
| Native SD | CNP | 0.241 ± 0.017 | 38.7 ± 3.1 | 6.479 ± 1.483 | 1.838 ± 0.283 | 0.262 ± 0.009 | 43.1 ± 2.2 | 6.719 ± 1.243 | 1.978 ± 0.221 |
| Deep Ensemble | ActFound | 0.377 ± — | 65.3 ± — | 75.246 ± — | 5.605 ± — | 0.392 ± — | 68.4 ± — | 61.236 ± — | 6.006 ± — |
| Deep Ensemble | MetaNN | 0.336 ± — | 56.4 ± — | 22.936 ± — | 3.435 ± — | 0.318 ± — | 53.4 ± — | 16.630 ± — | 2.874 ± — |
| MC Dropout | ActFound | 0.376 ± 0.001 | 65.1 ± 0.4 | 28.096 ± 0.767 | 5.132 ± 0.066 | 0.408 ± 0.002 | 72.0 ± 0.2 | 34.043 ± 0.995 | 6.144 ± 0.103 |
| MC Dropout | MetaNN | 0.354 ± 0.005 | 60.9 ± 1.0 | 14.548 ± 0.832 | 3.591 ± 0.128 | 0.351 ± 0.005 | 60.1 ± 1.0 | 12.577 ± 0.632 | 3.335 ± 0.109 |
| *clustering split* |  |  |  |  |  |  |  |  |  |
| Native SD | DKT | **0.158 ± 0.011** | **21.8 ± 2.8** | **1.876 ± 0.221** | **0.871 ± 0.098** | **0.201 ± 0.018** | **29.8 ± 4.0** | **2.462 ± 0.380** | **1.094 ± 0.142** |
| Native SD | ADKF | **0.111 ± 0.003** | **10.1 ± 0.4** | **1.175 ± 0.009** | **0.529 ± 0.005** | **0.096 ± 0.002** | **10.0 ± 0.6** | **1.220 ± 0.007** | **0.481 ± 0.005** |
| Native SD | CNP | 0.229 ± 0.016 | 36.7 ± 2.9 | 6.703 ± 1.392 | 1.829 ± 0.259 | 0.282 ± 0.009 | 47.5 ± 1.9 | 9.872 ± 1.730 | 2.439 ± 0.246 |
| Deep Ensemble | ActFound | 0.385 ± — | 67.7 ± — | 51.441 ± — | 5.951 ± — | 0.405 ± — | 70.4 ± — | 77.259 ± — | 7.190 ± — |
| Deep Ensemble | MetaNN | 0.339 ± — | 57.2 ± — | 24.747 ± — | 3.625 ± — | 0.336 ± — | 57.7 ± — | 16.857 ± — | 3.222 ± — |
| MC Dropout | ActFound | 0.398 ± 0.003 | 69.7 ± 0.7 | 44.542 ± 2.084 | 6.600 ± 0.101 | 0.427 ± 0.004 | 75.5 ± 0.6 | 62.636 ± 2.049 | 8.466 ± 0.150 |
| MC Dropout | MetaNN | 0.354 ± 0.005 | 60.5 ± 1.4 | 16.494 ± 1.178 | 3.832 ± 0.177 | 0.378 ± 0.006 | 65.5 ± 1.0 | 18.379 ± 1.208 | 4.216 ± 0.132 |

We quantified uncertainty for each query by the standard deviation (SD) of its predictive output. DKT, ADKF and CNP provide native SD during inference as the square root of their predictive variance. For ActFound and MetaNN, we estimated predictive SD using MC Dropout or Deep Ensemble. MC Dropout uses the population SD of predictions from 50 stochastic forward passes through one fitted model, with dropout active in each pass. Deep Ensemble uses the population SD across predictions from four independently trained models without dropout. The corresponding mean across passes or models provides the prediction evaluated with each estimator.

To assess robustness to training randomness, we repeated the native SD and MC Dropout evaluations across four independent training runs. Table 1 reports the mean and between-run sample SD of each calibration metric; this SD measures variation in evaluation results across runs. Deep Ensemble combines all four independently trained models into a single ensemble, leaving no independent ensemble replicates from which to estimate between-ensemble sample SD. Table 1 therefore shows a dash in place of this unavailable SD.

ADKF achieved the lowest mean MCA, CE90, NLL and ENCE in all four dataset–split combinations, followed by DKT and CNP (Table 1). This ordering was more consistent than the prediction-accuracy ordering in Figure 2: ActFound led on MAE under the clustering split, but even CNP had lower values on all four calibration metrics than either uncertainty estimator for ActFound or MetaNN. Thus, better prediction accuracy did not necessarily translate into better uncertainty calibration.

ADKF’s advantage over DKT persisted under the clustering split, although both GP models had higher MCA, CE90, NLL and ENCE than under the random split. On CHEMBL37, CE90 increased from 5.8 to 10.0 percentage points for ADKF and from 24.1 to 29.8 percentage points for DKT. ADKF also had smaller between-run sample SDs for all four metrics in every dataset–split combination, indicating more stable calibration across the four training runs. Targeted paired MCA comparisons supported ADKF’s advantage over DKT in all four combinations (Table 2C); other calibration comparisons remain descriptive.

**Table 2.** Statistical support for ICSR risk ranking, selective prediction and reconstruction, with selected secondary comparisons.

| A. ICSR versus matched risk scores |  |  |  |  |  |  |  |
| --- | --- | --- | --- | --- | --- | --- | --- |
| Panel | Model | Native SD |  |  | GP-DNR |  |  |
| | | nAURC | $\rho_{\text{risk}}$ | $R_{50}$ | nAURC | $\rho_{\text{risk}}$ | $R_{50}$ |
| *Random split* |  |  |  |  |  |  |  |
| ChEMBL27 | DKT | + | + | + | + | + | + |
| ChEMBL27 | ADKF | + | + | + | + | + | − |
| CHEMBL37 | DKT | + | + | + | + | + | + |
| CHEMBL37 | ADKF | + | + | + | − | + | − |
| *clustering split* |  |  |  |  |  |  |  |
| ChEMBL27 | DKT | + | + | − | − | − | − |
| ChEMBL27 | ADKF | + | + | − | − | − | − |
| CHEMBL37 | DKT | + | + | − | − | + | + |
| CHEMBL37 | ADKF | + | + | + | − | − | − |

| B. Core reconstruction tests: clustering split |  |  |  |  |
| --- | --- | --- | --- | --- |
| Diagnostic | ChEMBL27 |  | CHEMBL37 |  |
|  | DKT | ADKF | DKT | ADKF |
| Adjusted factor–error rank correlation ($\rho_{\text{adj}}$) | + | + | + | + |
| Factor-group normalized MAE gap difference | + | + | + | + |
| nAURC improvement over shuffled ICSR | + | + | + | + |

| C. Secondary comparisons: difference [95% interval] |  |  |  |  |
| --- | --- | --- | --- | --- |
| Comparison (metric) | ChEMBL27 |  | CHEMBL37 |  |
|  | Random split | clustering split | Random split | clustering split |
| ActFound vs DKT (MAE) | −0.0329<br>[−0.0492, −0.0181] | +0.0196<br>[0.0001, 0.0406] | −0.0134<br>[−0.0281, 0.0039] | +0.0114<br>[−0.0024, 0.0292] |
| ActFound vs ADKF (MAE) | −0.0212<br>[−0.0345, −0.0086] | +0.0170<br>[−0.0025, 0.0389] | −0.0093<br>[−0.0203, 0.0055] | +0.0154<br>[0.0025, 0.0332] |
| ADKF vs DKT (MCA) | +0.0380<br>[0.0252, 0.0530] | +0.0465<br>[0.0301, 0.0653] | +0.0888<br>[0.0686, 0.1147] | +0.1041<br>[0.0886, 0.1252] |

MetaNN outperformed ActFound on all four calibration metrics with both MC Dropout and Deep Ensemble, despite its poorer prediction accuracy. Changing the uncertainty estimator had a less consistent effect than changing the predictor. Compared with MC Dropout, Deep Ensemble reduced MetaNN’s MCA, CE90 and ENCE in every dataset–split combination, but increased its NLL in three of the four; CHEMBL37 under the clustering split was the exception. For ActFound, Deep Ensemble increased NLL in all four combinations, while its effects on the other metrics depended on the dataset and split. Neither estimator closed the gap to DKT or ADKF.

ADKF achieved mean observed interval coverage closest to the nominal 90% level in all four dataset–split combinations, whereas DKT’s and CNP’s intervals contained substantially fewer query activities (Figure 3). On CHEMBL37, ADKF’s mean observed interval coverage at this level was 90.1% under the random split and 84.6% under the clustering split, compared with 66.4% and 60.6% for DKT. The curves also exposed residual miscalibration in ADKF: observed interval coverage exceeded nominal interval coverage at intermediate levels on ChEMBL27 under the random split, but fell below it at high nominal levels on both datasets under the clustering split. Agreement in the mean curve can also mask discrepancies within individual assays.

**Figure 3.**
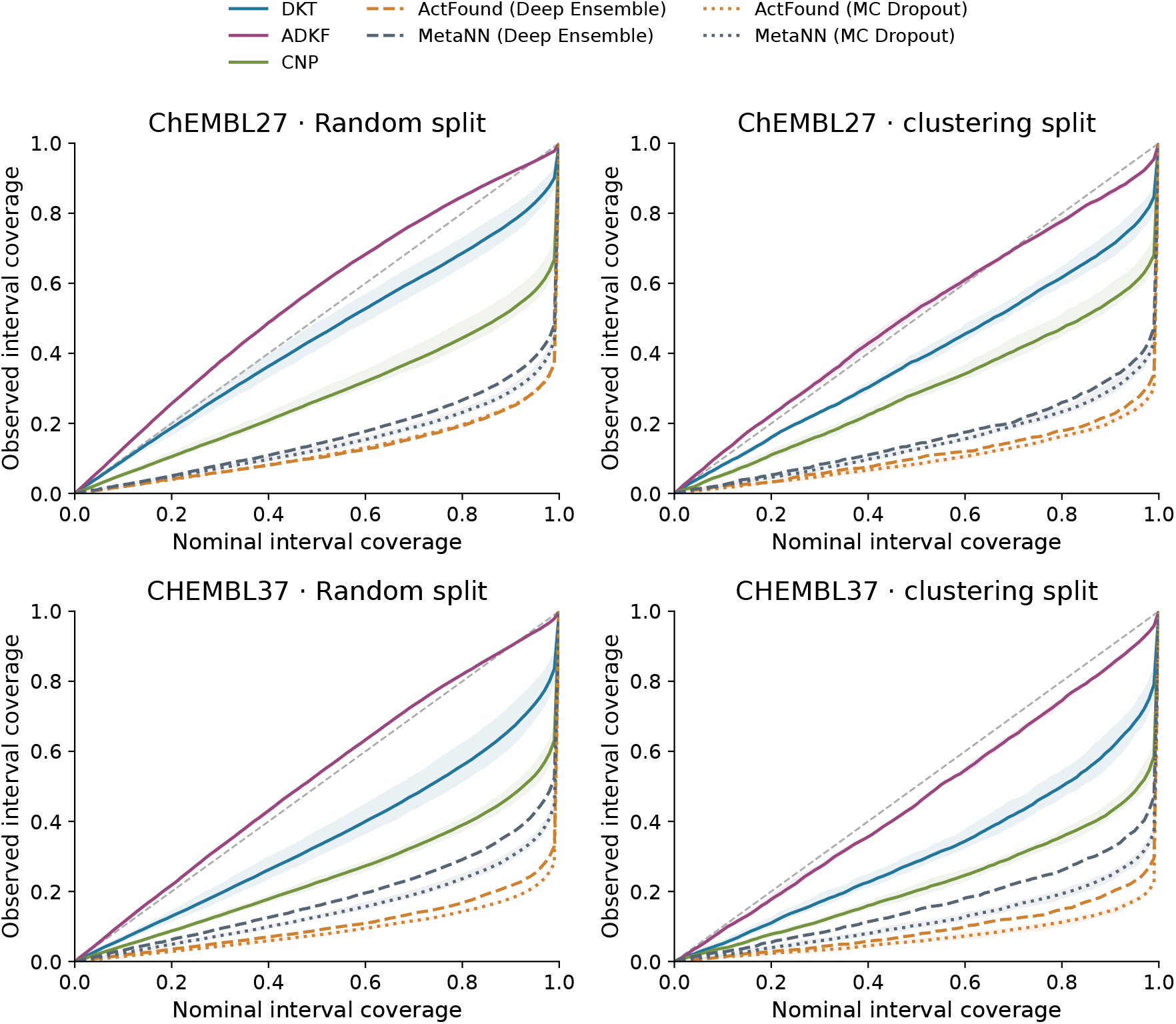
Observed versus nominal interval coverage for native, Deep Ensemble and MC Dropout uncertainty. Nominal interval coverage is the target probability assigned to each prediction interval. Observed interval coverage is the fraction of all query activities contained in their corresponding intervals within a support–query split. Solid lines denote native SD, dashed colored lines Deep Ensemble, and dotted lines MC Dropout. Shaded regions span the observed range across four independent training runs for native SD and MC Dropout; each Deep Ensemble curve describes one four-member ensemble and has no range band. Observed interval coverage is averaged over support–query splits within each assay and then equally over 100 assays. The gray diagonal indicates agreement between observed and nominal interval coverage; shading is not a confidence interval.

MetaNN’s curves lay above ActFound’s through most nominal levels with both uncertainty estimators, yet both remained far below the diagonal at high nominal levels (Figure 3). On CHEMBL37 under the clustering split, for example, MetaNN’s mean observed interval coverage at the nominal 90% level was 32.3% with Deep Ensemble and 24.5% with MC Dropout. This improvement was small relative to the remaining discrepancy between observed and nominal interval coverage.

MetaNN’s closer agreement with nominal interval coverage coincided with broader predictive distributions than ActFound’s. Averaged over assays, its mean, median and 90th-percentile predictive SDs were larger with both estimators in every dataset and split, despite higher RMSE. Broader distributions may partly explain why its intervals contained more query activities; this comparison does not isolate that effect from how uncertainty was assigned across queries. The lower MCA and CE90 should therefore be interpreted alongside prediction accuracy and interval sharpness, the concentration of the predictive distribution.^22^ The comparison also involves different variance components: DKT and ADKF include observation noise in their predictive variances, whereas Deep Ensemble and MC Dropout omit an observation-noise term.

### 2.3 Risk ranking and selective prediction with DKT and ADKF

In this section, we compared risk ranking with native SD, GP-DNR and ICSR for DKT and ADKF using three metrics: risk–error Spearman correlation (*ρ*_risk_), normalized area under the risk–coverage curve (nAURC) and half-query MAE reduction (*R*_50_). *ρ*_risk_ is the Pearson correlation between the ranks of risk scores and the ranks of absolute query prediction errors; higher values indicate better agreement with the actual error ordering. nAURC averages MAE relative to full-query MAE across successively smaller sets of the lowest-risk queries; lower values indicate better selective prediction. *R*_50_ measures the percentage MAE reduction achieved by retaining the lowest-risk half; higher values indicate greater error reduction. Figure 4 summarizes overall performance, while Figures 5 and 6 show the error reduction achieved by retaining progressively lower-risk queries.

**Figure 4.**
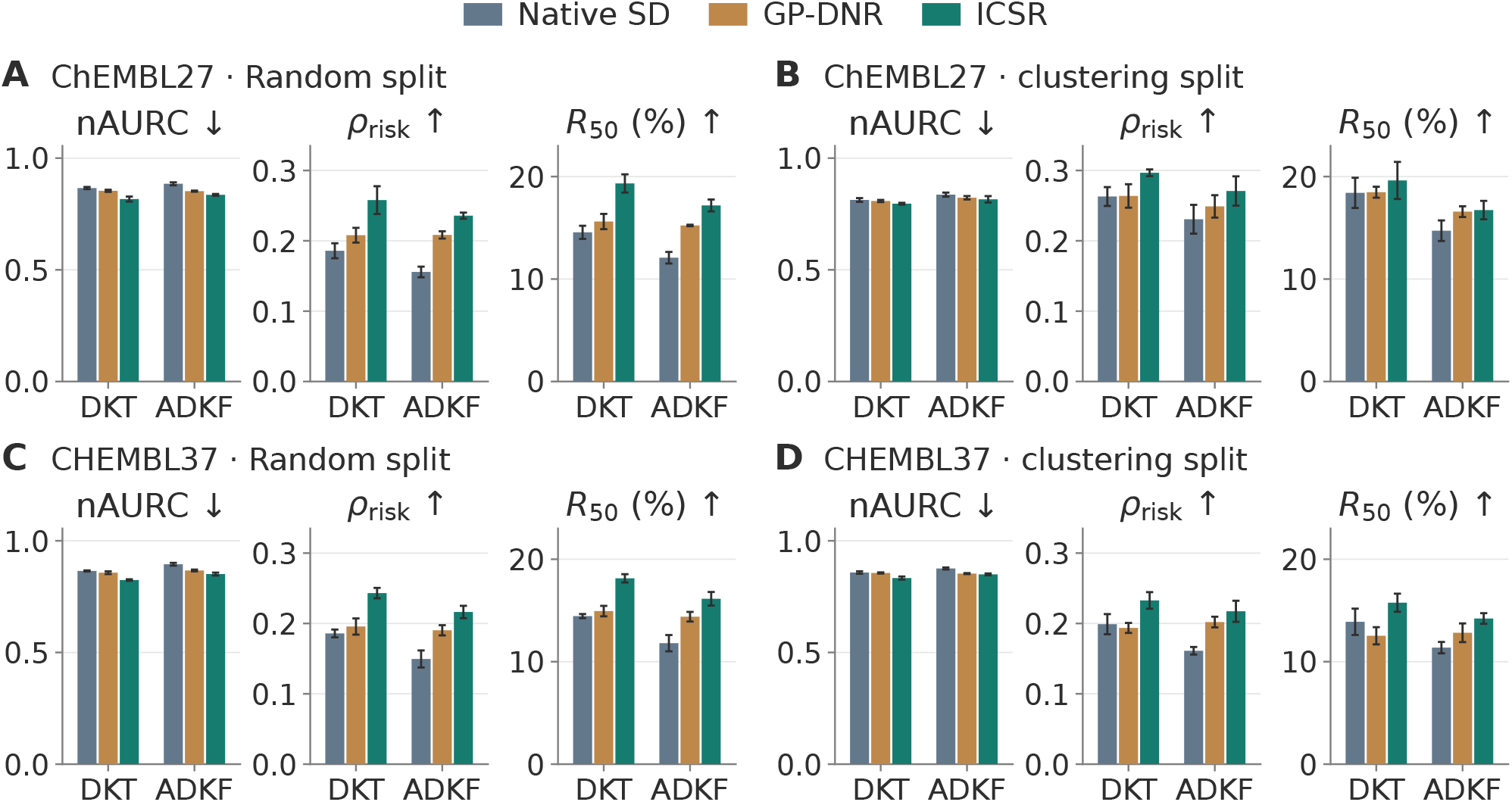
Risk ranking with native SD, GP-DNR and ICSR for DKT and ADKF. Normalized area under the risk– coverage curve (nAURC) averages normalized retained-query MAE over all retained counts; the denominator is full-query MAE from the same fitted model and support–query split, and coverage denotes query retention fraction. Risk– error Spearman correlation (*ρ*_risk_) correlates risk ranks with absolute-error ranks. Half-query MAE reduction (*R*_50_) is 100[1 − MAE_ret_ (⌈*N*_*Q*_/2⌉)/MAE_full_], the percentage MAE reduction for the lowest-risk half, rounded up. Lower nAURC and higher *ρ*_risk_ and *R*_50_ are favorable. Panels show ChEMBL27 under the random split (A) and clustering split (B), and CHEMBL37 under the random split (C) and clustering split (D), with 100 assays per dataset. Bars show means, and error bars show between-run sample SDs across four independent training runs, after random-repeat averaging within assay and equal assay weighting. Scores share predictions within each fitted model; comparisons between backbones are descriptive. ICSR uses validation-selected *η* = 0.75 for both backbones and *κ* = 4 for DKT and 16 for ADKF. Each metric has its own scale, fixed across panels; all bar axes start at zero.

**Figure 5.**
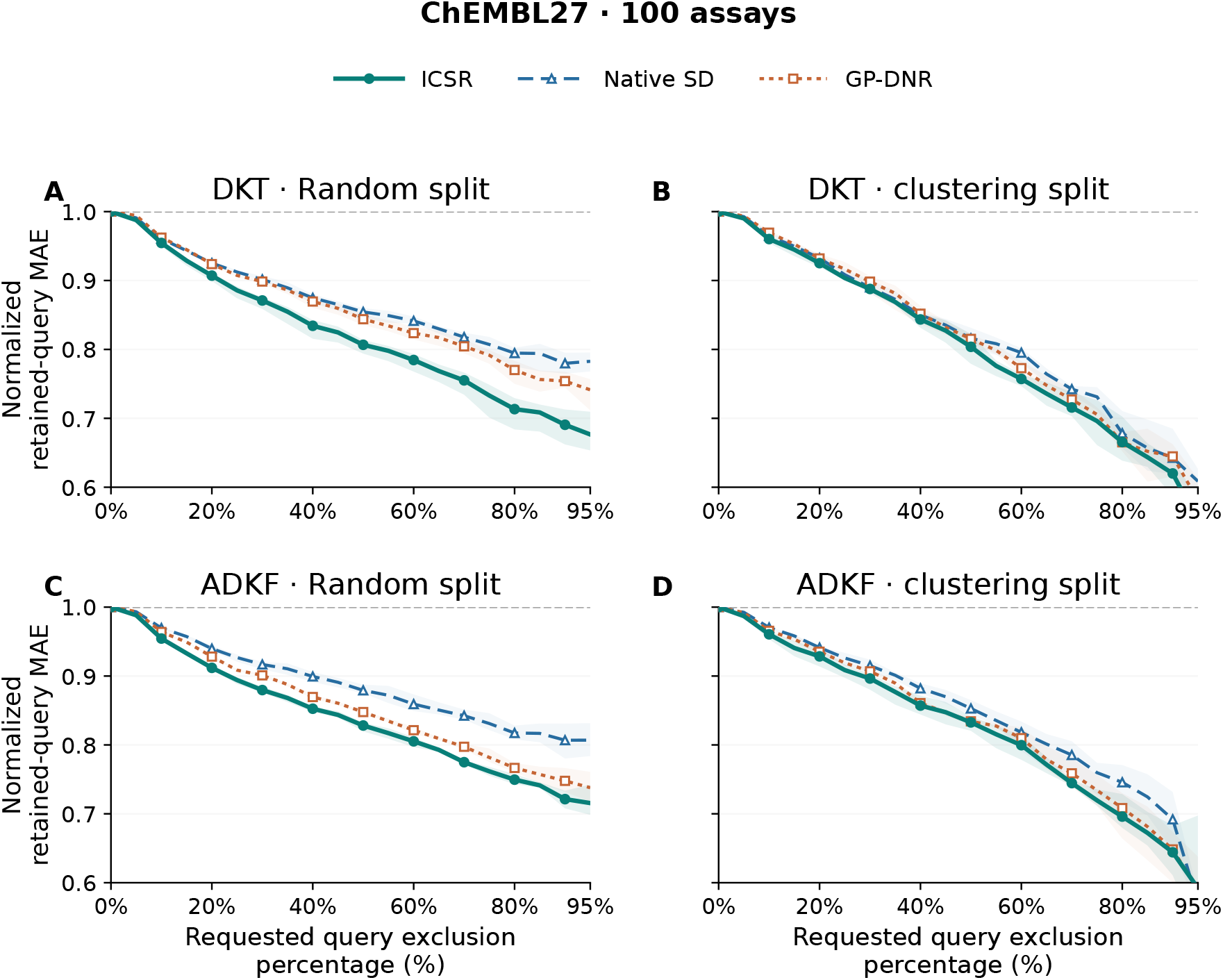
Normalized retained-query MAE in 100 ChEMBL27 test assays. Normalized retained-query MAE is MAE among the retained lowest-risk queries divided by full-query MAE for the same fitted model and support–query split; one denotes unchanged MAE and lower values indicate lower error. DKT occupies the first row (A,B) and ADKF the second (C,D); columns show the random split and the clustering split. At requested query exclusion percentage 100*u*, the lowest-risk ⌈(1 − *u*) *N*_*Q*_⌉ queries remain. All curves start at one, and the displayed range is 0.6–1.0. Requested query exclusion percentage runs from 0 to 95% in five-percentage-point steps; retained counts and normalization denominators match across scores. Colors and line styles identify ICSR (solid circles), native SD (dashed triangles) and GP-DNR (dotted squares). Ratios average random repeats within assay, then 100 assays equally, and finally metrics from four independent training runs. Bands show the observed range across runs, not confidence intervals. Segments join observed values without smoothing or a monotonic fit.

**Figure 6.**
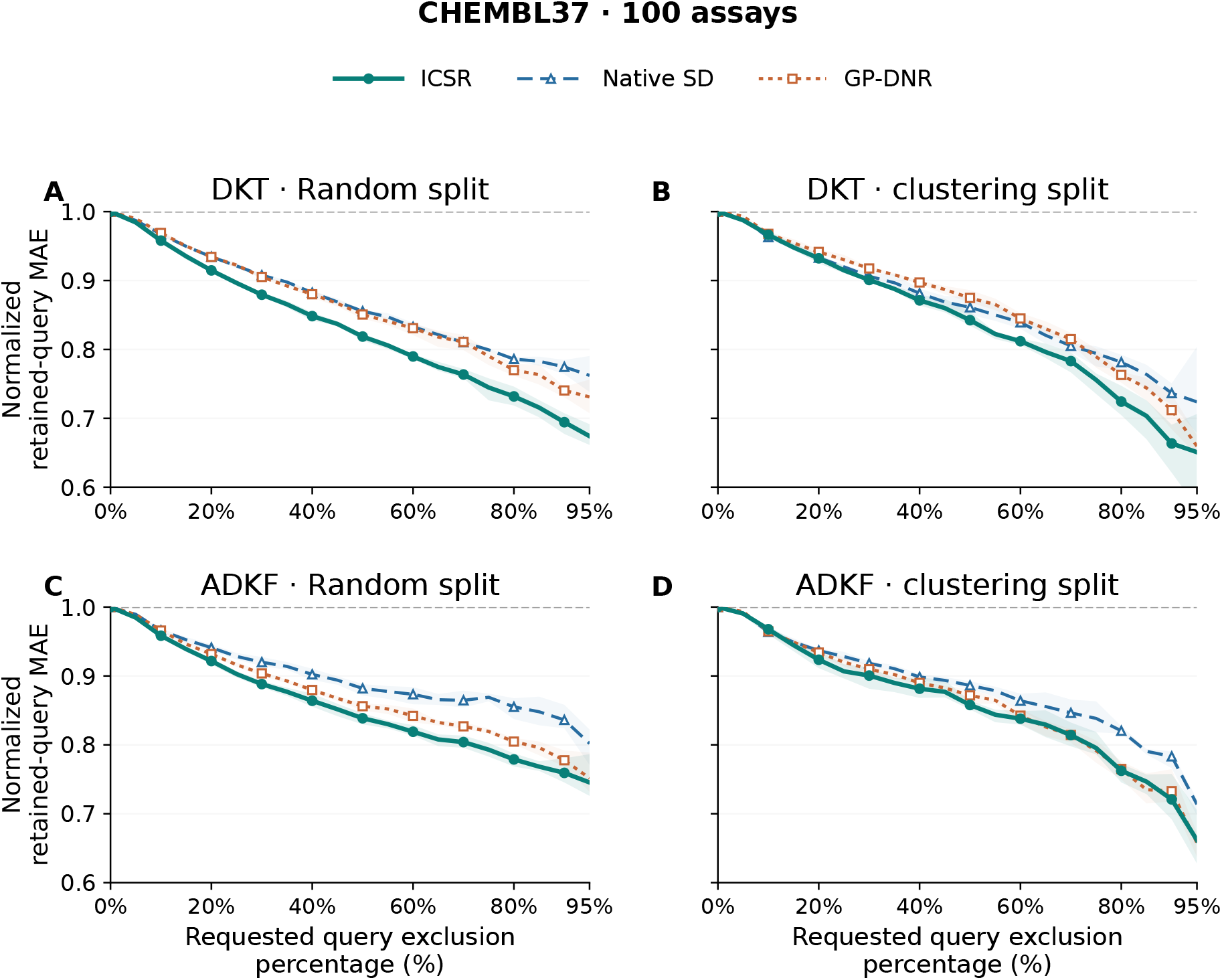
Normalized retained-query MAE in 100 CHEMBL37 assays. Normalized retained-query MAE is MAE among the retained lowest-risk queries divided by full-query MAE for the same fitted model and support–query split; lower values indicate lower error. The first row shows DKT (A,B) and the second ADKF (C,D), with the random split on the left and the clustering split on the right. Each panel compares ICSR, native SD and GP-DNR as the requested query exclusion percentage increases. At requested query exclusion percentage 100*u*, each score retains the lowest-risk ⌈(1 − *u*) *N*_*Q*_⌉ queries from the same predictions. All curves start at one. Retained counts, equal-assay aggregation, observed ranges across runs, colors and axis limits match Figure 5.

Under the random split, ICSR achieved the lowest mean nAURC and highest mean *ρ*_risk_ and *R*_50_ with both backbones on both datasets (Figure 4A,C). GP-DNR also improved on native SD, but ICSR provided a larger additional gain for DKT than for ADKF. DKT with ICSR gave the best mean performance among the six model–score combinations: its *R*_50_ was 19.3% on ChEMBL27 and 18.1% on CHEMBL37, compared with 17.2% and 16.1% for ADKF with ICSR. Within each fitted model, predictions were shared across scores, so these gains reflect better identification of lower-error queries.

Under the clustering split, ICSR retained the best means for each backbone, and DKT with ICSR again led all six combinations (Figure 4B,D). ICSR’s nAURC advantage over GP-DNR was smaller than under the random split for both backbones. GP-DNR provided little gain over DKT’s native SD on ChEMBL27 and had lower *ρ*_risk_ and *R*_50_ on CHEMBL37, whereas ICSR improved all three metrics. With ADKF, GP-DNR captured much of the gain over native SD; *R*_50_ on ChEMBL27 was 16.6% with GP-DNR and 16.7% with ICSR. The additional gain over GP-DNR was therefore clearer for DKT than for ADKF under this split.

Removing the highest-risk queries progressively reduced normalized retained-query MAE under the random split on both datasets (Figures 5A,C and 6A,C). ICSR produced the lowest mean curve at every nonzero displayed query exclusion percentage for both backbones. Its curves continued to fall as selection became stricter, whereas native SD flattened or rose slightly at some late points. At 90% requested query exclusion, DKT with ICSR reached normalized retained-query MAE of 0.69 on both datasets, compared with 0.72 on ChEMBL27 and 0.76 on CHEMBL37 for ADKF with ICSR. Retaining lower-risk queries therefore produced sustained relative error reduction with ICSR, with larger reductions for DKT at this selection level.

The clustering split also showed lower normalized retained-query MAE as progressively lower-risk queries were retained, although the separation between scores varied with the selection level (Figures 5B,D and 6B,D). For DKT, ICSR maintained a lower curve than GP-DNR at every nonzero displayed query exclusion percentage on CHEMBL37; on ChEMBL27 it nearly coincided with GP-DNR at 80% requested query exclusion. For ADKF, ICSR and GP-DNR followed similar curves and converged under strict selection. At 95% requested query exclusion, ADKF had slightly lower normalized retained-query MAE with GP-DNR than with ICSR on both datasets, and its native SD gave the lowest value on ChEMBL27. Lower-risk selection therefore reduced error overall, but ICSR’s better mean performance in Figure 4 did not imply the lowest error at every selection level.

### 2.4 Support reconstruction failure carries information about query error

The reconstruction factor ( *f*_*q*_) for query *q* is a dimensionless multiplier of native predictive variance, built from errors in predicting measured supports from the other supports. The detailed formulation and rationale are given in Methods (Section 4.3; eq 3). It weights squared reconstruction errors, standardized by reconstruction variance, by each support’s influence on a query—its contribution to the fitted prediction’s sensitivity—and stabilizes the weighted average toward the overall support mean. A factor above one increases the native variance; a factor below one decreases it. ICSR equals native SD multiplied by the square root of the reconstruction factor. This factor is distinct from the evaluation correlations *ρ*_risk_ and *ρ*_adj_. Two tests under the clustering split examined whether the reconstruction factor contained additional query-error information and whether its usefulness depended on which supports carried the reconstruction errors.

First, we tested whether larger reconstruction factors were associated with larger absolute prediction errors for query compounds after controlling for two other quantities. Native predictive variance is the variance of the model’s predictive distribution for a query compound. Maximum query–support Tanimoto similarity is the highest Tanimoto similarity between a query compound and any measured support, using binary Morgan fingerprints; higher values indicate a closer structural analogue. Both quantities may themselves be associated with absolute prediction error, so controlling for them tests whether the reconstruction factor provides additional information about absolute prediction error.

Within each assay, we first converted the reconstruction factor, absolute prediction error, native predictive variance and maximum query–support Tanimoto similarity to ranks. We then fitted two linear regressions, using the ranks of native predictive variance and maximum query–support Tanimoto similarity to predict reconstruction factor ranks and absolute prediction error ranks separately. We subtracted the fitted values from the corresponding reconstruction factor ranks and absolute prediction error ranks, then correlated the two sets of residuals. This gives the partial rank correlation (*ρ*_adj_); a positive value indicates that larger reconstruction factors remain associated with larger absolute prediction errors after accounting for these linear relationships on the rank scale. Across all queries, mean *ρ*_adj_ was 0.14 for DKT and 0.12 for ADKF on ChEMBL27, and 0.09 and 0.13 on CHEMBL37; all four intervals excluded zero (Table 2B).

We also examined whether the reconstruction factor distinguished absolute prediction error among queries with relatively close measured analogues. Within each assay, we selected the third of queries with the highest maximum query–support Tanimoto similarity and divided them into equal groups with lower and higher reconstruction factors. We compared normalized group MAE, defined as the mean absolute prediction error within each group divided by full-query MAE from the same fitted model and support–query split. The denominator includes all queries, including those outside the high-similarity subset; higher normalized group MAE indicates larger absolute prediction errors. On ChEMBL27, normalized group MAE increased from 0.70 to 0.91 for DKT and from 0.70 to 0.97 for ADKF between the groups with lower and higher reconstruction factors (Figure 7A,C). The reconstruction factor therefore distinguished absolute prediction error even among queries with relatively close measured analogues.

**Figure 7.**
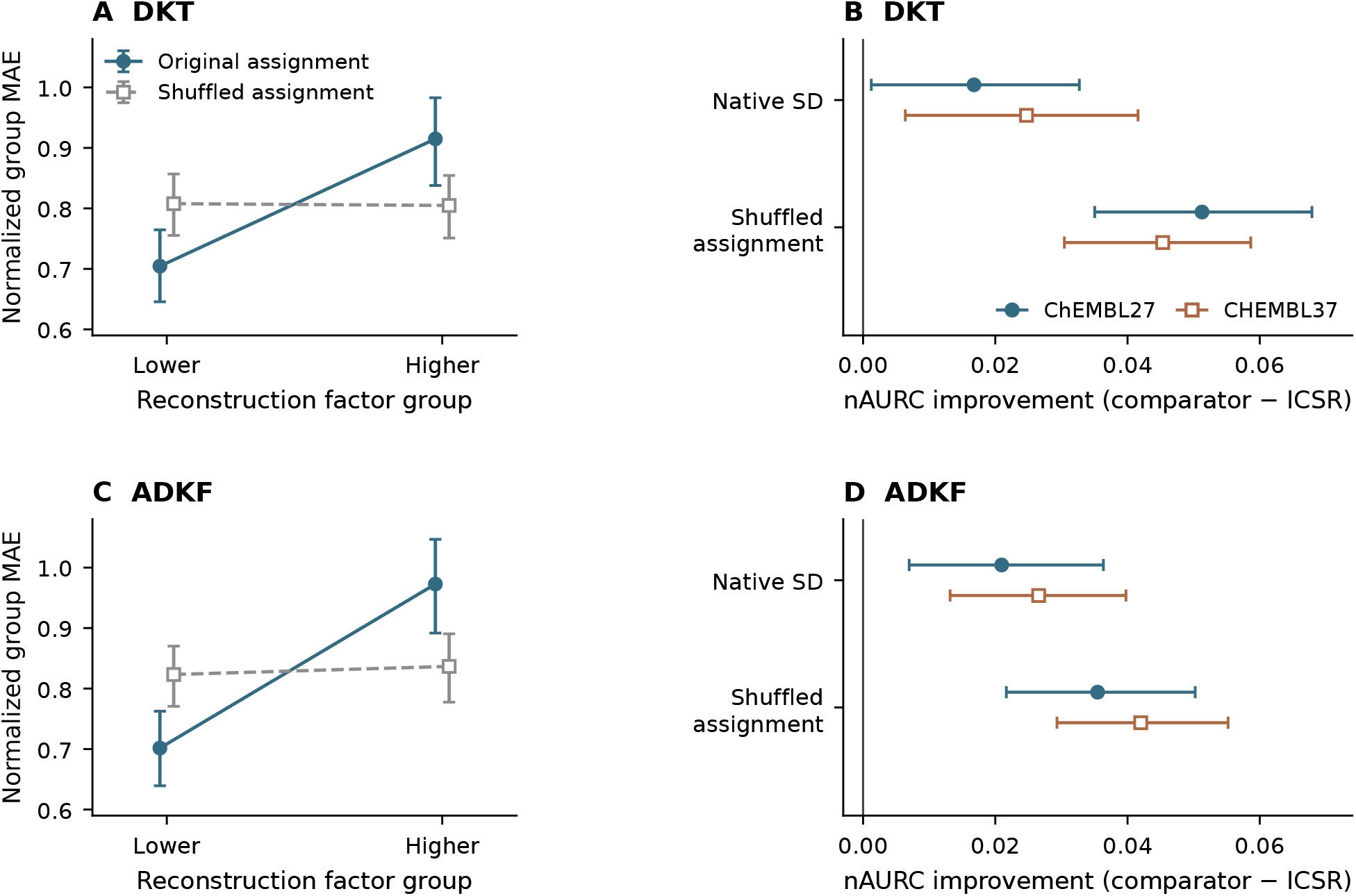
Normalized group MAE and nAURC improvement assess reconstruction information in ICSR. Normalized group MAE is MAE within a reconstruction-factor group divided by full-query MAE; higher values indicate larger group errors. The nAURC improvement is comparator nAURC minus ICSR nAURC; positive differences favor ICSR. nAURC averages normalized retained-query MAE over every retained query count. Supports are measured compounds used by the model; queries have held-out activities for evaluation. Reconstruction predicts each support activity from the other supports with the fitted model fixed. The reconstruction factor combines squared reconstruction errors, standardized by reconstruction variance, using query-specific influence and stabilization; it multiplies native predictive variance. (A,C) ChEMBL27 queries in each assay’s highest third of maximum query–support Tanimoto similarity (binary Morgan feature presence) are divided into equal lower and upper reconstruction-factor groups. An odd central position is omitted; exact ties use expected ordering. Shuffling reassigns the same errors among supports and recomputes groups. (B,D) nAURC improvements on all queries. Native SD is the model’s predictive SD. Shuffled ICSR preserves variance, weights and stabilization. Both panels contain 100 assays evaluated under the clustering split. Means weight assays equally before averaging four training-seed metrics. Bars show 95% source-group bootstrap intervals from 5,000 draws preserving all fitted models; differences are paired. Shuffled results average 20 permutations.

Second, we tested whether information from support reconstruction contributed to ICSR’s assessment of prediction risk for query compounds. If knowing which supports reconstructed poorly provided no useful information about query prediction risk, randomly reassigning support reconstruction errors among supports would be expected to leave risk-ranking performance largely unchanged. We therefore shuffled the already computed standardized squared support reconstruction errors among supports 20 times. This preserved the set of standardized squared support reconstruction errors while disrupting their correspondence with support identities. We recomputed reconstruction factors and ICSR risk scores after each shuffle, keeping influence weights, support selection, stabilization, native predictive variance and query activity predictions fixed.

Shuffling reduced ICSR’s ability to prioritize query predictions with smaller absolute prediction errors. Across all queries, nAURC increased by 0.035–0.051 after shuffling across the two backbones and datasets (Figure 7B,D). A higher nAURC indicates larger normalized retained-query MAE averaged over all query retention counts. All four paired intervals supported this loss of performance. Original ICSR also had nAURC 0.017–0.027 lower than native SD, with all four paired intervals supporting improvement. The high-similarity query subset showed a corresponding loss of absolute prediction error separation. The normalized group MAE difference between the groups with higher and lower reconstruction factors (higher minus lower) fell from 0.21 to approximately zero for DKT and from 0.27 to 0.01 for ADKF on ChEMBL27, with the same loss of separation on CHEMBL37 (Figure 7A,C). These results support the contribution of support reconstruction information, correctly assigned to individual supports, to ICSR’s assessment of query prediction risk.

### 2.5 Statistical significance

Statistical testing focused on ICSR versus native SD and GP-DNR within each backbone on nAURC, *ρ*_risk_ and *R*_50_, and on reconstruction-factor associations and shuffled-assignment controls. Selected comparisons also assessed ActFound versus each GP backbone on MAE and ADKF versus DKT on MCA (Table 2). We computed 95% percentile bootstrap intervals by resampling connected assay-source groups, retaining all fitted-model results for each sampled assay and pairing comparisons by assay and split. Intervals excluding zero indicated significance. These intervals were conditional on the fitted models and unadjusted for multiple comparisons; other comparisons and the ASAP application remained descriptive.

ICSR significantly improved nAURC and *ρ*_risk_ over native SD in all eight dataset–split–backbone combinations. Its *R*_50_ gains were significant in all random-split settings and, under the clustering split, only for ADKF on CHEMBL37. Gains over GP-DNR were more often significant under the random split; under the clustering split, no nAURC gain was significant, and only DKT on CHEMBL37 significantly improved *ρ*_risk_ and *R*_50_ (Table 2A). All four clustering-split settings showed significantly positive *ρ*_adj_ and significantly larger factor-group normalized MAE gaps and lower nAURC with original than shuffled assignments (Table 2B). ADKF had significantly lower MCA than DKT in all four dataset–split combinations. Significant MAE differences favored both GP models under the random split on ChEMBL27, and ActFound under the clustering split against DKT on ChEMBL27 and ADKF on CHEMBL37; the remaining MAE comparisons were inconclusive (Table 2C).

### 2.6 Retrospective application to SARS-CoV-2 Mpro potency prediction

Maximum query–support Tanimoto similarity (*T*_max_) is the largest binary Morgan fingerprint Tanimoto similarity between a query and any measured support; its mean across queries describes proximity to available structural analogues. Half-query MAE reduction (*R*_50_) measures the percentage decrease in MAE after retaining the lowest-risk half. We retrospectively selected the SARS-CoV-2 Mpro assay for its favorable *R*_50_ to illustrate selective prediction within a chemical series containing close query–support structural analogues (Figure 8). The released Train/Test memberships from the public ASAP–Polaris–OpenADMET challenge supplied 842 support and 263 query measurements, with mean *T*_max_ of 0.803.^23,24^

**Figure 8.**
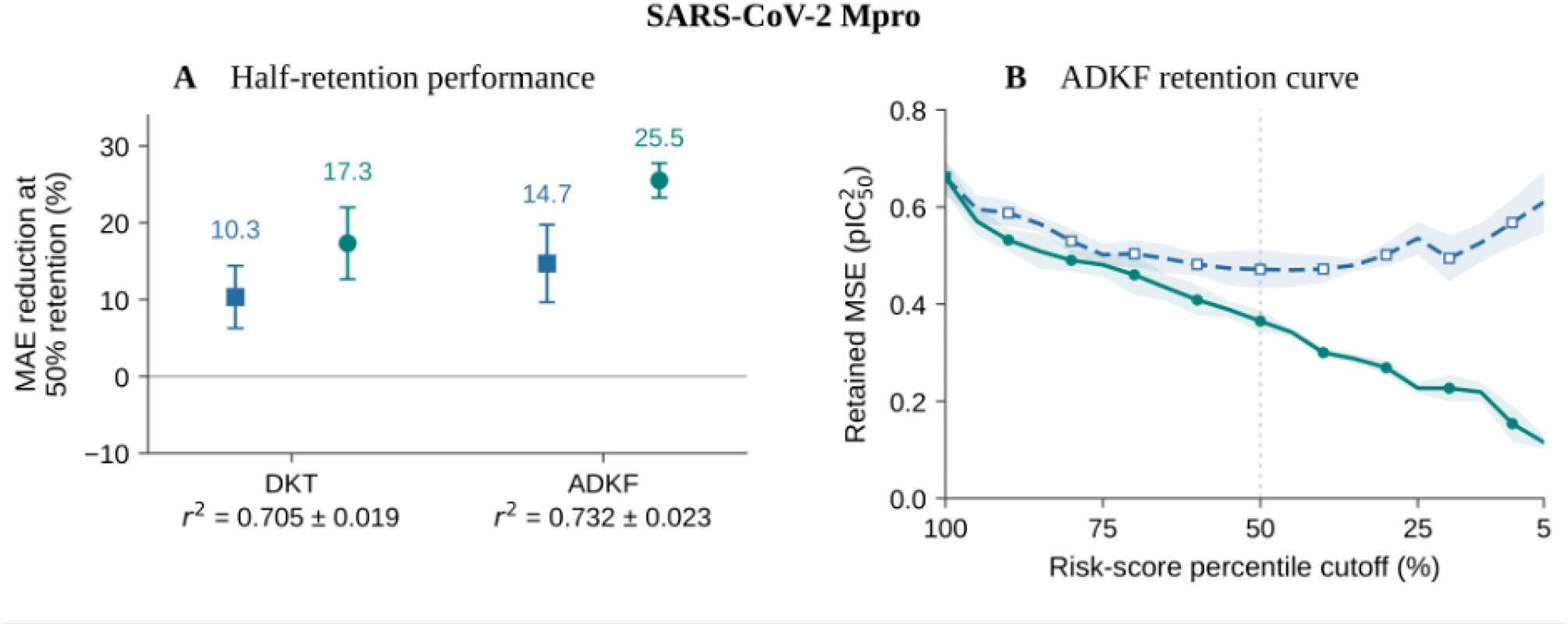
Half-query MAE reduction (*R*_50_), squared Pearson correlation (*r*^2^) and retained-query MSE in the retrospectively selected ASAP SARS-CoV-2 Mpro case (842 supports; 263 queries). *R*_50_ is the percentage decrease in MAE after retaining the lowest-risk half, rounded up; higher values are favorable. *r*^2^ is the squared correlation between predicted and measured activities, with higher values indicating stronger unsigned linear association. Retained-query MSE is mean squared prediction error among retained queries, in squared pIC_50_ units; lower values are favorable. Blue squares denote native SD; green circles denote ICSR. (A) *R*_50_ for DKT and ADKF. Filled symbols show four-run means with between-run sample SD whiskers. Values beneath each backbone give full-query *r*^2^ (mean ± between-run sample SD). (B) ADKF retained-query MSE as the risk-score percentile cutoff decreases from 100 to 5%, in 5-percentage-point steps. A cutoff of *p* corresponds to a requested query retention percentage of *p*: each score retains the lowest-risk ⌈*pN*_*Q*_/100⌉ queries. Lines show means across the four runs and shading denotes between-run sample SD, not confidence intervals. The vertical line marks the 50% cutoff used for *R*_50_ (132/263 queries). Predictions and retained counts are identical between scores within each run. Points are joined without smoothing or monotonic fitting. ADKF illustrates the backbone with the larger observed *R*_50_ improvement in this assay.

Squared Pearson correlation (*r*^2^) is the square of the correlation between predicted and measured query activities; higher values indicate stronger unsigned linear association. Between-run sample SD describes variation in *r*^2^ across the four training runs. Before risk-based selection, full-query *r*^2^ was 0.705 ± 0.019 for DKT and 0.732 ± 0.023 for ADKF (means ± between-run sample SDs; Figure 8A). These full-query *r*^2^ values are shared by native SD and ICSR because risk scoring preserves the predictions.

Half-query MAE reduction (*R*_50_) is the percentage decrease in average absolute prediction error after retaining the lowest-risk ⌈*N*_*Q*_/2⌉ queries, relative to all *N*_*Q*_ queries; higher *R*_50_ is favorable. ICSR improved mean *R*_50_ from 10.3 to 17.3% for DKT and from 14.7 to 25.5% for ADKF (Figure 8A). ADKF’s 10.8-percentage-point improvement was the larger of the two mean differences: retaining the ICSR-selected half reduced MAE by approximately one-quarter relative to the full query set.

Retained-query MSE is mean squared error (MSE) among the retained lowest-risk queries, expressed in squared pIC_50_ units; lower values indicate smaller squared prediction errors. At requested query retention percentage *p*, each score retains ⌈*pN*_*Q*_/100⌉ queries, so the actual query retention fraction can slightly exceed *p*/100. For ADKF, mean retained-query MSE decreased as the requested query retention percentage decreased (Figure 8B). It fell from 0.661 over all queries to 0.365 for the ICSR-selected half, compared with 0.471 for native SD, and reached 0.116 at 5% requested query retention. Within each fitted model, both scores used identical predictions and retained counts throughout the curves. The lower retained-query MSE therefore reflects selection of more reliable predictions from the existing query set.

## 3. Discussion and Conclusions

### 3.1 Uncertainty benchmarking in multi-task bioactivity prediction

The benchmark identifies different strengths across predictor–uncertainty combinations in multi-task bioactivity prediction. Under shared support–query splits, ActFound, DKT and ADKF provided the strongest mean point predictions, while ADKF led the calibration diagnostics. DKT’s native SD ranked errors more effectively than ADKF’s (Figure 2; Table 1; Figure 4). The best mean point predictions also depended on the split, with the GP backbones favored by the random split and ActFound by the clustering split. These findings support assessing prediction accuracy, calibration and risk ranking together when selecting a predictor and its uncertainty estimator. An overall change in uncertainty scale can bring observed interval coverage closer to its nominal level while leaving the ordering of queries unchanged, so calibration alone does not establish the ability to identify reliable predictions.

MetaNN’s lower MCA, CE90, NLL and ENCE than ActFound, despite less accurate predictions and broader uncertainty scales, further illustrate the need to consider calibration alongside sharpness.^22^ The ensemble and dropout estimates here omit observation noise, whereas native model variances include it. Their calibration comparison consequently concerns the specific uncertainty formulations evaluated. Assessing each estimator against its own errors describes the complete predictor–uncertainty combination; matched comparisons within a fitted model isolate the contribution of the risk score.

### 3.2 Contribution of ICSR to risk assessment

ICSR improves the identification of reliable bioactivity predictions by connecting measured support reconstruction errors to individual queries. Across both GP backbones, assay panels and split regimes, it improved all three mean risk-ranking metrics over native SD and the adapted GP-DNR comparator (Figure 4). DKT with ICSR achieved the best mean ranking metrics among the six DKT/ADKF model–score combinations. Because the matched scores use identical activity predictions, these gains reflect better selection from an existing predictor. GP-DNR contributes information about local activity variation; ICSR contributes evidence of how well the fitted model explains its measured supports.

The reconstruction controls support the value of this model-specific evidence. Adjusted factor–error rank correlation (*ρ*_adj_) remained modestly positive after controlling for maximum query–support Tanimoto similarity and native predictive variance. Positive factor-group normalized MAE gaps also showed error separation among close analogues. Shuffling the same reconstruction errors among supports reduced both the factor-group normalized MAE gap and risk-ranking performance (Figure 7; Table 2B). Their usefulness therefore depends on where errors occur as well as their magnitude. Influence weighting connects those errors to queries through the fitted GP’s dependence on supports. Stabilization reduces reliance on a few dominant errors by drawing on the full-support mean, extending the use of standardized LOO errors for global GP variance-scale estimation to query-specific risk assessment.^18^

### 3.3 Query–support similarity and selective prediction

The relationship between supports and queries shapes the selection problem. The random split can place close chemical analogues in both sets, while the clustering split tests transfer between chemical groups. The resulting changes in accuracy and calibration emphasize the need to evaluate uncertainty under the intended chemical conditions, consistent with molecular benchmarks comparing random, scaffold and chronological splits.^25^ The clustering split increased the GP backbones’ MAE even when normalized retained-query MAE improved. Because nAURC averages normalized retained-query MAE within each support–query split, practical evaluation should consider retained-query MAE on the assay scale and the intended query retention fraction as well as relative ranking performance (Figures 5 and 6).

The retrospectively selected SARS-CoV-2 Mpro case illustrates selective prediction within a series containing close query–support structural analogues. ADKF’s half-query MAE reduction (*R*_50_) increased from 14.7% with native SD to 25.5% with ICSR, and retained-query MSE decreased as ICSR selection became stricter (Figure 8). This setting allows measured analogues to inform which existing predictions are more reliable. Such an ordering could support compound assessment by identifying estimates suitable for initial use and flagging higher-risk predictions for additional measurement.

### 3.4 Limitations and future validation

The main applicability question is how well support reconstruction informs queries with low structural similarity to their closest support. ICSR’s nAURC improvements over native SD and GP-DNR were smaller under the clustering split than under the random split, and the paired intervals for the nAURC comparisons with GP-DNR under the clustering split included zero (Table 2A). This pattern suggests that greater query– support structural similarity may favor the transfer of reconstruction information to queries. Direct tests varying support–query similarity and support density are needed to establish this dependence. Retention curves also show that the preferred score can change for very small retained subsets, making the intended selection threshold part of the application conditions.

ICSR also inherits the learned representation and covariance assumptions of its GP backbone. Reconstruction with a fixed fit assesses agreement with measured supports and may miss errors that arise only in unrepresented chemistry. The present evidence covers DKT and ADKF. For each backbone, retention and stabilization were selected on the 76 ChEMBL27 validation assays and then held fixed across all test conditions; transfer to other backbones and assay settings requires further assessment.

Predictive intervals derived from ICSR would require separate validation of interval coverage. LOO residuals can be correlated,^26^ so reconstruction-based rescaling alone does not establish agreement between observed and nominal interval coverage. Conformal regression for QSAR^27^ and jackknife+^28^ offer interval constructions with interval coverage guarantees under their respective assumptions. Combining reconstruction information with such constructions remains a direction for future work. Prospective evaluations should further test whether selecting predictions with ICSR improves experimental hit rates or reduces measurement costs.

This study provides a systematic benchmark of uncertainty estimation in multi-task bioactivity prediction and introduces ICSR for improving query risk ranking. The benchmark supports evaluating accuracy, calibration and error ordering as complementary objectives; ICSR adds measured reconstruction evidence to select more reliable predictions from fitted GP models. Query–support structural similarity and the intended query retention fraction guide its use, while prospective validation is needed to establish its value for experimental decisions.

## 4. Methods

### 4.1 Data

We use ChEMBL27 as the name for the public 4,276-assay collection reported by Martin et al. in 2019 and distributed in the ActFound data release.^1,29^ This name identifies the dataset used here; its original ChEMBL release has not been independently verified. We adopted the MetaMix Group 1 assay split, with 4,100 assays for training, 76 for validation and 100 for internal testing.^30^

The CHEMBL37 dataset came from ChEMBL release 37 (2026), released after the original study.^31^ We applied the original authors’ public extraction and preparation pipeline, including activity conversion, structure processing and within-assay duplicate handling, retaining assays with at least 50 compounds and an activity SD of at least 0.5.^32^ After excluding all 4,276 ChEMBL27 assay identities, we sampled 100 assays uniformly without replacement from 4,261 eligible assays. The resulting CHEMBL37 test set contains 14,495 assay–compound records and shares no assay identities with ChEMBL27.

The retrospective application used the SARS-CoV-2 main protease (Mpro) potency dataset from the public ASAP–Polaris–OpenADMET version 2 release, preserving all 842 Train and 263 Test measurements as supports and queries, respectively.^23,24^

### 4.2 Predictors and uncertainty estimators

We implemented DKT, ADKF and a conditional neural process (CNP) from scratch, using the same two-layer fingerprint encoder architecture with separately trained weights.^12,13,33^ DKT conditions a globally learned GP head on assay supports; ADKF adapts its GP head to those supports. Both use a Matérn covariance. CNP aggregates support representations and activities into a context for its query decoder. ActFound and MetaNN were trained using the ActFound authors’ source code.^2^

Uncertainty quantification used either native model outputs or variability across repeated predictions. DKT and ADKF provide predictive variances that include observation noise, and CNP provides variance through its Gaussian decoder. The square root of each variance defines native SD.

For calibration of ActFound and MetaNN, Deep Ensemble combined four models trained independently without dropout, using the mean and population SD of their predictions as the ensemble prediction and uncertainty.^34^ MC Dropout models were trained separately with dropout probability 0.1 and evaluated with dropout active for 50 stochastic forward passes per checkpoint; the draw mean and population SD supplied that checkpoint’s prediction and uncertainty.^35^ Neither estimator added an observation-noise term.

### 4.3 Influence-weighted reconstruction

ICSR combines support reconstruction scores with query-specific influence weights and stabilizes their weighted average toward an assay-wide reference. For support set *S*, define 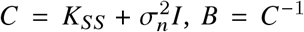 and *α* = *B*y_*S*_ in support-standardized coordinates, where *K*_*SS*_ is the fitted support kernel matrix and 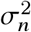 is observation-noise variance. The fitted encoder, GP parameters and full-support label transform remain fixed throughout reconstruction; ADKF’s head is adapted once before these calculations.

For each support *s*, we reconstruct its activity from the remaining supports. Let 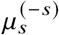 and 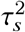 denote the corresponding conditional mean and variance. The reconstruction score is the squared error relative to this variance:

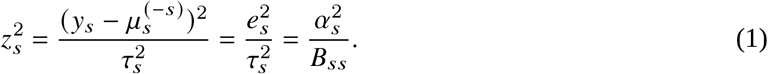

Here 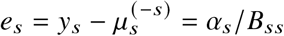 and 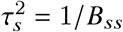 follow the standard analytic GP leave-one-out identities.^14^ Variance-normalized squared leave-one-out errors have previously been used to estimate a global GP variance scale.^18^ We use them as support-level reconstruction scores: a larger value indicates poorer agreement with the measured activity relative to the model’s conditional uncertainty.

We define influence as the expected squared change in the fitted query prediction after removing a support. For query *q* with fitted mean *μ*_*q*_ = *k*_*qS*_ *B*y_*S*_, sensitivity to support activity is *h*_*qs*_ = ∂ *μ*_*q*_/∂ *y*_*s*_ = [*k*_*qS*_ *B*]_*s*_. Block inversion gives 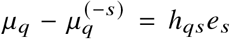. Under the fixed GP reference 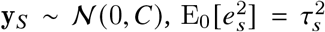, yielding

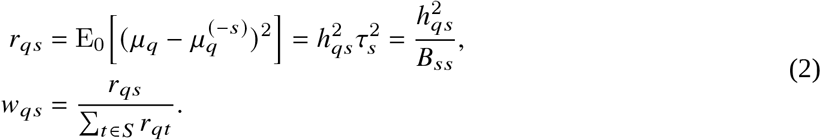

Thus, influence is defined by expected changes in the fitted prediction, with *B* accounting for covariance among supports; the weights are not normalized pairwise kernel similarities. The identity 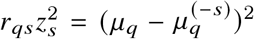 connects the reconstruction score to the observed deletion effect on each query. The Gaussian reference motivates the weighting and does not assert empirical calibration.

For each query, retain the ⌈*η*|*S*|⌉ highest-weight supports in *T*_*q*_, resolving exact ties by stable record identifiers, and renormalize their weights to 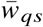. Their weighted reconstruction score is 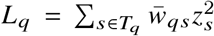. Influence weighting can concentrate this score on a few reconstruction errors. We therefore construct a stabilization rule that retains local information while drawing on the full-support mean 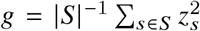. Using *g* as the reference preserves the assay’s observed reconstruction-error scale when local information is limited:

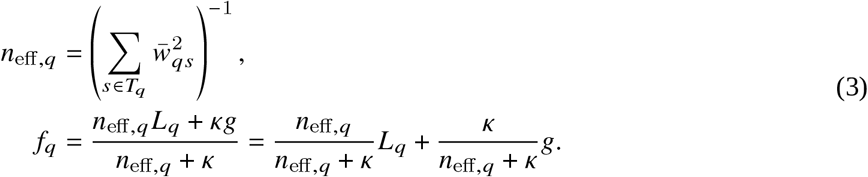

The nonnegative parameter *κ* controls the strength of stabilization. The effective count equals *m* for *m* equally weighted supports and approaches one when a single support dominates. Using this count instead of |*T*_*q*_ | makes stabilization respond to how broadly influence is distributed. At fixed *κ* > 0, concentrated weights increase the global contribution *κ*/(*n*_eff,*q*_ + *κ*), whereas distributed weights preserve more local information. Here *n*_eff,*q*_ measures weight concentration; it does not assume independent reconstruction errors.

The reconstruction factor *f*_*q*_ is a query-specific variance multiplier, distinct from the risk–error Spearman correlation *ρ*_risk_ and adjusted factor–error rank correlation *ρ*_adj_ used for evaluation. The nonnegative coefficients in this blend sum to one, keeping *f*_*q*_ between *L*_*q*_ and *g*. Setting *κ* = 0 recovers the local score, while increasing *κ* without bound approaches the global score. When *L*_*q*_ = *g* = 1, the multiplier remains one. These properties motivate our construction as a gradual, query-dependent adjustment of the balance between local and assay-wide reconstruction evidence. The final risk is 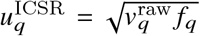, where 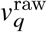 is native predictive variance on the recorded activity scale. Predicted activities and conditioning on all supports remain unchanged.

We selected (*η, κ*) for each backbone on the 76 ChEMBL27 validation assays by minimum nAURC under the clustering split, using *η* ∈ {0.25, 0.5, 0.75, 1} and *κ* ∈ {0, 0.5, 1, 2, 4, 8, 16, 32, 64, ∞}. Metrics were averaged over support–query splits within each assay, assays received equal weight and metrics were averaged over four training seeds; exact ties favored larger *η*, then smaller *κ*. The selected settings were *η* = 0.75 for both backbones, *κ* = 4 for DKT and *κ* = 16 for ADKF, fixed before recomputing both test panels and shared across fitted models and split regimes.

### 4.4 Risk baselines

Risk ranking and selective prediction were evaluated for DKT and ADKF by comparing ICSR with native SD, 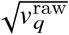, and an adapted GP-DNR baseline.

Kötter et al.’s GP-DNR supplements GP uncertainty with predicted local structure–activity variation. Using 2048-bit Morgan fingerprints of radius 2, they define neighbors by Tanimoto similarity greater than 0.5 and the different-neighbor ratio (DNR) as the fraction of neighbors whose activities differ by at least one log unit. An auxiliary GP learns DNR from training compounds with neighbors; compounds without neighbors are excluded. Its predicted DNR is added to the activity GP’s uncertainty. The original GP models use Tanimoto kernels in FlowMO.^20^

We independently reimplemented the published procedure for DKT and ADKF, retaining its neighborhood rules, auxiliary DNR prediction and additive construction. For each episode, we fitted an auxiliary Tanimoto GP in scikit-learn to supports with neighbors, using one recorded activity unit for the activity-gap threshold. We clipped predicted DNR to [0, 1] and added it to the fitted backbone’s raw predictive variance, giving the risk score 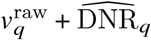. This adaptation combines the backbone’s uncertainty with local activity variation while preserving identical activity predictions for the native SD, GP-DNR and ICSR comparisons.

### 4.5 Metrics

Prediction accuracy was assessed using mean absolute error (MAE) and squared Pearson correlation (*r*^2^). Root mean squared error (RMSE) supports the calibration diagnostics, while mean squared error (MSE) is used for the application selection curves. These error metrics use query errors *e*_*q*_ = *y*_*q*_ − *ŷ*_*q*_ on the recorded activity scale. For all *N*_*Q*_ queries in one support–query split,

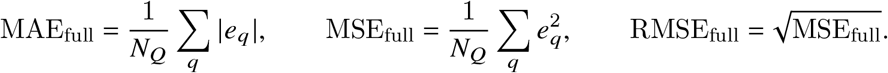

Lower values indicate smaller errors. MAE and RMSE have activity units; MSE has squared activity units. Squared Pearson correlation is *r*^2^ = corr(*y, ŷ*)^2^, computed within each split before averaging. It measures unsigned linear association and differs from the residual-based coefficient of determination *R*^2^.

Nominal interval coverage is the target probability *p* assigned to a prediction interval; observed interval coverage is the fraction of all query activities contained in their corresponding intervals. We assessed each uncertainty estimator against its own prediction errors. For a central Gaussian interval based on prediction *ŷ*_*q*_ and evaluated predictive SD 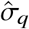, observed interval coverage in one support–query split is

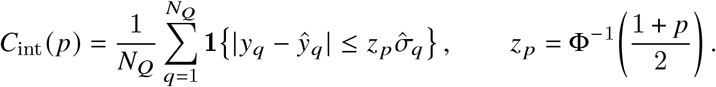

Here 1{·} is the indicator function and Φ is the standard normal cumulative distribution function. Miscalibration area (MCA) integrates the absolute discrepancy |*C*_int_( *p*) − *p*| across the evaluated nominal interval coverage levels. Absolute interval coverage error at 90% (CE90) is 100 |*C*_int_(0.90) −0.90| percentage points, computed within each split before averaging. Gaussian negative log likelihood (NLL) is the mean negative natural logarithm of the Gaussian predictive density at measured activities:

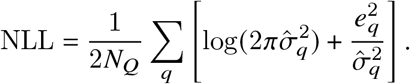

NLL depends on both prediction error and predictive SD, and on the recorded activity scale. Expected normalized calibration error (ENCE) averages |RMSE−RMV|/RMV across ten approximately equal-count bins ordered by predicted variance. Root mean predicted variance (RMV) is the square root of the mean predicted variance within a bin; RMSE is computed from the same queries.^36^ Lower MCA, CE90, NLL and ENCE are favorable. Predictive SD describes uncertainty for one query; its mean, median and 10th and 90th percentiles summarize the distribution of uncertainty width within an assay.

Risk–error Spearman correlation (*ρ*_risk_) is the Pearson correlation between the ranks of risk scores and absolute errors; higher values indicate better ordering of queries by error magnitude. The query retention fraction is *k*/*N*_*Q*_ after retaining the *k* lowest-risk queries. Let MAE_ret_(*k*) be MAE within this subset. Normalized retained-query MAE divides this quantity by full-query MAE. Normalized area under the risk–coverage curve (nAURC) and half-query MAE reduction (*R*_50_) are

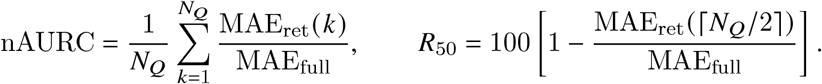

Lower nAURC and higher *R*_50_ are favorable. *R*_50_ is a percentage reduction within one split; differences in *R*_50_ between scores are percentage points. Exact risk ties use expected random ordering, and matched scores share predictions and retained counts. The requested query exclusion percentage 100*u* in the benchmark curves retains ⌈(1 − *u*) *N*_*Q*_⌉ queries. The requested query retention percentage *p* in the application retains ⌈*pN*_*Q*_/100⌉ queries. These requested percentages specify the count before rounding; the actual query retention fraction is always *k*/*N*_*Q*_. Retained-query MSE is the mean squared error within the retained subset, reported in squared pIC_50_ units for the application.

Adjusted factor–error rank correlation (*ρ*_adj_) measures the within-assay partial rank association between the reconstruction factor and absolute query error after controlling for maximum query–support Tanimoto similarity and native predictive variance. We regress each of the first two rank vectors on an intercept and the two control rank vectors, then correlate the residuals. Positive *ρ*_adj_ indicates remaining factor–error association after adjustment. Normalized group MAE divides a reconstruction-factor group’s MAE by full-query MAE from the same fitted model and split. The factor-group normalized MAE gap is upper-factor normalized group MAE minus lower-factor normalized group MAE. Its original-minus-shuffled difference tests how reconstruction-error assignment affects separation. The nAURC improvement is comparator minus ICSR, whereas improvements in *ρ*_risk_ and *R*_50_ are ICSR minus comparator; positive differences favor ICSR.

Between-run sample SD describes variation in a metric across four independent training runs, using the sample denominator of three; it differs from the predictive SD assigned to individual queries. Metrics were computed per fitted model or ensemble within each support–query split, then averaged over random repeats within assay and equally across assays. Single-model summaries report means and between-run sample SDs; each Deep Ensemble provides one estimate without independent ensemble replicates. Paired comparisons use unadjusted 95% source-group bootstrap intervals, with comparison families and draw counts specified in Results. Secondary comparison families were selected after inspecting mean results.

## Data and software availability

The accompanying single-model package contains manuscript sources, tables, figures, model-level metrics, paired comparisons and reproducible analysis scripts. A source manifest identifies the frozen model arrays and data records used by the analysis. ChEMBL27 and CHEMBL37 are available through the ActFound Figshare release and ChEMBL, respectively, under their upstream terms.^29,31^ The application data and published Train/Test memberships are available from the ASAP Zenodo record.^24^ The repository records check-point identities, downloaded-source hashes and inference provenance. A permanent public archive identifier and final software license have not yet been assigned.

## Author information

**Corresponding author.** Li Tian.

## Author contributions

Z.W.: Conceptualization, Methodology, Software, Validation, Formal analysis, Investigation, Data curation, Visualization, Writing—original draft, and Writing—review and editing. L.T.: Conceptualization, Supervision, Project administration, and Writing—review and editing. C.-M.C.: Supervision, Resources, Project administration, Funding acquisition, and Writing—review and editing.

## Acknowledgments

This work was supported by the Laboratory for Synthetic Chemistry and Chemical Biology under the Health@InnoHK Program launched by the Innovation and Technology Commission, the Government of the Hong Kong Special Administrative Region of the People’s Republic of China. Z.W. acknowledges the Laboratory for Synthetic Chemistry and Chemical Biology for a studentship from 2022 to 2025.

